# Effects of early life stress paired with adolescent alcohol consumption on two-bottle choice alcohol drinking behaviors in mice

**DOI:** 10.1101/2023.01.21.524642

**Authors:** Thomas W. Perry, Harrison M. Carvour, Amanda N. Reichert, Elizabeth A. Sneddon, Charlotte A.E.G. Roemer, Ying Ying Gao, Kristen M. Schuh, Natalie A. Shand, Jennifer J. Quinn, Anna K. Radke

## Abstract

**Background:** In humans, early life stress (ELS) is associated with an increased risk for developing both alcohol use disorder (AUD) and post-traumatic stress disorder (PTSD). We previously used an infant footshock model that produces stress-enhanced fear learning (SEFL) in rats and mice and increases aversion-resistant alcohol drinking in rats to explore this shared predisposition. The goal of the current study was to extend this model of comorbid PTSD and AUD to male and female C57BL/6J mice.

**Methods:** Acute ELS was induced using 15 footshocks on postnatal day (PND) 17. In adulthood, alcohol drinking behavior was tested in one of three two-bottle choice drinking paradigms. In continuous access, mice were given 24 h access to 5% or 10% ethanol and water for five consecutive drinking sessions each. In limited access drinking in the dark, mice were given 2 h of access to 15% ethanol and water across 15 sessions 3 h into the dark cycle. In intermittent access, mice were presented with 20% ethanol and water Monday, Wednesday, and Friday, for four consecutive weeks. In a fifth week of intermittent access drinking, increasing concentrations of quinine (10 mg/L, 100 mg/L, and 200 mg/L) were added to the ethanol to test aversion-resistant drinking. Intermittent access drinking was tested with and without a period of adolescent drinking (PND 35).

**Results:** Infant footshock did not alter drinking in the continuous or limited access tasks. Adult consumption and preference were lower in the intermittent access task when adolescent drinking was included and there were ELS-induced differences in consumption of quinine-adulterated ethanol in females.

**Conclusions:** Our results demonstrate that infant footshock followed by a period of adolescent drinking is a viable model of comorbid PTSD and AUD in rats and mice.

## Introduction

Alcohol use disorder (AUD) is a complex psychiatric disorder influenced by both genetic and environmental factors (Mayfield et al., 2008; Tawa et al., 2016; Young-Wolff et al., 2011). Stress is one environmental variable that is strongly associated with alcohol abuse and the development of AUD (Guinle and Sinha, 2020; Sinha, 2001). Among individuals with PTSD, AUD is reported to be up to three times more prevalent than it is in the general public (Gilpin and Weiner, 2017; Kessler et al., 1995; Smith and Cottler, 2018). This suggests that PTSD and AUD may be products of common alterations brought about by stress. Early life stress (ELS) in humans is associated with an increased risk of alcohol dependence, early onset alcohol use, and PTSD-AUD comorbidity (Enoch, 2011; Lee et al., 2018; Pechtel and Pizzagalli, 2011). These associations indicate that in early stages of development the brain may be particularly susceptible to stress-induced changes (Lupien et al., 2009).

Consistent with clinical literature, in preclinical models, chronic ELS in the form of isolation or maternal separation generally increases levels of alcohol consumption later in adulthood (Becker et al., 2011; Cruz et al., 2008; McCool and Chappell, 2009). Additionally, a number of studies have found that ELS increases aversion-resistant drinking when alcohol is paired with an aversive stimulus such as quinine (Bertagna et al., 2021; Radke et al., 2020; Shaw et al., 2020). This drinking despite a negative consequence has been used to model compulsive-like drinking, a central characteristic of AUD (Hopf et al., 2010; Hopf and Lesscher, 2014; Radke et al., 2017; Sneddon et al., 2019). Sex has been reported to play a role in the relationship between ELS and alcohol-drinking behaviors; however, the exact influence of sex remains unclear (Bertagna et al., 2021; Radke et al., 2020; Shaw et al., 2020).

In a recent study using Long-Evans rats, we found that acute ELS in the form of repeated footshock on postnatal day (PND) 17 increased aversion-resistant drinking in an intermittent access alcohol drinking paradigm (Radke et al., 2020). This form of acute infantile trauma is commonly used to sensitize fear learning, a phenomenon known as stress-enhanced fear learning (SEFL), as a preclinical model for PTSD and PTSD susceptibility (e.g., Poulos et al., 2014; Quinn et al., 2014; Minshall et al., in press). The common influence of infant footshock on disordered drinking behavior and fear learning, but not other forms of hippocampal-dependent learning or cognitive flexibility (Sneddon et al., 2021), suggests that an acute infant footshock paradigm may be a viable stress model for investigating AUD- PTSD comorbidity, as well as the shared neural mechanisms underlying both disorders.

To extend the utility of this preclinical model, the current study explored how acute ELS in the form of infant footshock influences consumption of alcohol alone and when paired with an aversive outcome in C57BL/6J mice. We used an infant stress protocol (15 footshocks on PND 17) that we have previously found increases fear learning and resistance to extinction in mice (Sneddon et al., 2021). To comprehensively characterize the effects of infant footshock on drinking, adult mice were exposed to alcohol in one of three two-bottle choice drinking tasks: limited access drinking in the dark, continuous access, or intermittent access. Based on previous findings in rats that had undergone PND 17 footshock and alcohol drinking (Radke et al., 2020), we ran a second intermittent access task where mice were presented with alcohol during adolescence (PND 35-55) to measure the necessity of alcohol exposure during that developmental stage on adult outcomes. To model aversion-resistant drinking, alcohol was adulterated with increasing concentrations of quinine, an aversive bittering agent, in the drinking in the dark and intermittent access paradigms. Additionally, considering published findings (Hennessy et al., 2009) and data from our own labs (Reichert et al., unpublished data) showing that exposure to conspecifics following trauma, otherwise known as social buffering, can modulate the effect of trauma, we investigated whether social buffering could alter the effects of infantile trauma on intermittent access drinking. Finally, due to robust evidence of enhanced vulnerability to both stress-related disorders and addictive behaviors in females (Bangasser and Valentino, 2014; Minshall et al., 2024; Quigley et al., 2021; Radke et al., 2021b), we examined sex as a potential moderator of the effects of infant footshock on alcohol drinking in all experiments.

## Materials and Methods

### Subjects

C57BL/6J mice were generated from breeding pairs from The Jackson Laboratory (Bar Harbor, ME, USA). Mice were provided food (Rodent Diet 5001, Cincinnati Lab Supply, Cincinnati, OH) and reverse osmosis (RO) water *ad libitum*. Mice were on a 12:12 light/dark cycle. Other than PND 17 footshock and adolescent alcohol exposure, all behavioral tests were conducted during adulthood (PND 60+). During infancy, all mice were briefly anesthetized with isoflurane and given an ear snip for identification purposes. Prior to home cage drinking, mice were group housed (2-4 mice/cage). One week before the first drinking session mice were individually housed in standard shoe box Udel polysulfone rectangular mouse cages (18.4 cm x 29.2 cm x 12.7 cm) outfitted with 2-bottle cage tops. All animals were cared for in accordance with the guidelines set by the National Institute for Health. All procedures were approved by the Institutional Animal Care and Use Committee (IACUC) at Miami University.

### Early life stress exposure

Mice were placed in a novel MED-Associates conditioning chamber (Context A) on PND 17 (after Quinn et al., 2014). Context A was brightly lit, contained a uniform grid floor, was scented with vanilla, and was cleaned with odorless 5% sodium hydroxide. Mice received either 15 footshocks (1 mA, 1 sec) or no footshocks during a 60-min session beginning at 60 sec following placement in the chamber. Shocks were delivered according to a variable inter-shock interval of 120-240 sec. Progressive scan video cameras containing visible light filters (VID-CAMMONO-2A; Med Associates, Inc.) monitored mice throughout the session.

For the continuous access and drinking in the dark experiments, mice were returned to a cage with their littermates in a brightly lit room separate from the conditioning chambers following shock exposure. Mice were run in squads of 4 and the entire litter was returned to the home cage 3 h after the final squad’s session.

For the first intermittent access experiment (Experiment 3), we discovered an issue with the shock delivery boxes that resulted in animals receiving a variable number of shocks on PND 17. After determining that 6 or more footshocks produced the SEFL effect (i.e., significantly higher fear learning in adult mice exposed to shock vs. no shock) (Reichert et al., unpublished data), we excluded mice that received fewer than 6 footshocks from the drinking analyses. In this experiment, we also examined the effects of social buffering on SEFL (Reichert et al., unpublished data) and alcohol drinking in adulthood. Immediately following the ELS exposure session, mice were placed into their assigned social buffering condition for 90 minutes. During this time, pups in the isolation condition were housed individually while pups in the littermates condition were housed as a group. All buffering took place in polysulfone holding cages (19 x 29.2 x 12.7 cm) that are similar to the housing cages but without bedding. All holding cages were in a room with overhead lighting and were placed on top of Sunbeam-brand heated mats, half on the pad and half off. For the second intermittent access experiment (Experiment 4), all mice were in the isolation condition following footshock.

### Fear Conditioning and Memory Test

Before the commencement of drinking, on PND 60, mice that took part in intermittent access drinking (Experiments 3 and 4) were fear conditioned in a novel context. This context consisted of four identical conditioning chambers (32.4 x 25.4 x 21.6 cm; Med-Associates, Inc., Georgia, VT) housed inside sound-attenuating cabinets. The chambers were distinct from the PND 17 footshock and isolation contexts in several ways: the room was dark except lit by red lamps, the room smelled of vinegar, there were black Plexiglass triangular inserts within the chamber, and the floor had staggered stainless steel rods (18 rods, two rows, .05 cm vertically apart). Mice were placed into the fear conditioning chamber and received one footshock (0.25 mA, 1s) 180 seconds following placement within the chamber. Thirty seconds following footshock, mice were returned to the home cage. For Experiment 3, mice were administered a fear retention test session in this context on PND 90 to test incubation of fear (data not shown).

### Two-bottle choice ethanol drinking

For all experiments, mice were given access to RO drinking water and ethanol in RO water (v/v) in their home cage. Mice were weighed at the beginning of every session and solutions were made fresh for each drinking session. Bottles were alternated each session to equate side biases. Two “dummy” cages were outfitted with bottles to account for spillage and evaporation.

#### Experiment 1: Continuous access ethanol drinking

Forty mice (20 male and 20 female) were given access to 5% or 10% ethanol (**Figure 1A**). Each ethanol concentration was provided for 5 consecutive, 24-h continuous drinking sessions and bottle weights were recorded every 24 h.

**Figure 1.**
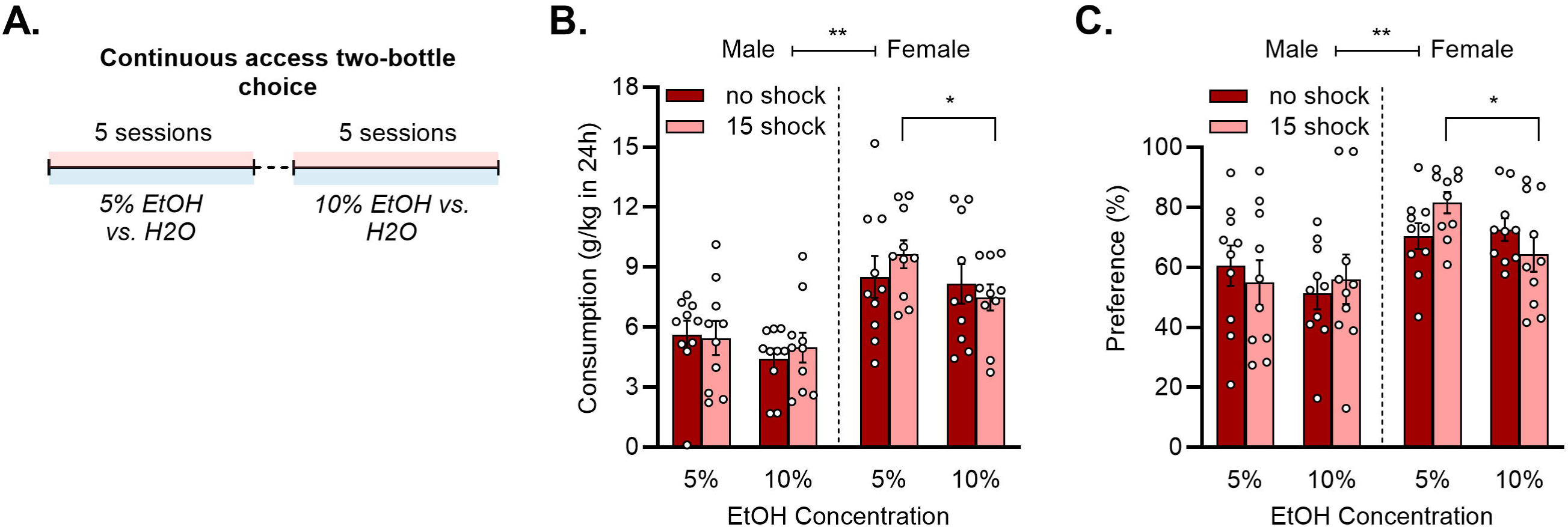
Sex, but not infant footshock, influences ethanol consumption and preference in a continuous access drinking task. (**A**) Mice (n = 10/group) drank 5% and 10% ethanol for five 24-h sessions each. (**B**) Infant footshock did not impact ethanol consumption (g/kg). Females consumed significantly more ethanol than males, **p < 0.001 main effect of sex. Female mice that received footshock consumed more 5% ethanol than 10% ethanol, *p = 0.008 Holm-Sidak’s test. (**C**) Infant footshock did not impact preference for ethanol vs. water. Ethanol preference was higher in females vs. males, **p = 0.002 main effect of sex. Ethanol preference was higher in female mice that received footshock at the 5% vs. 10% ethanol concentration, *p = 0.047 Holm-Sidak’s test.

#### Experiment 2: Limited access drinking in the dark

Fifty-five mice (28 male and 27 female) were presented with 15% ethanol following a limited access, two-bottle choice protocol for 2 h a day, five day/week (Monday-Friday) (**Figure 2A**). Bottles were placed on the home cage 3 h after lights out and bottle weights were recorded before and after each session.

**Figure 2.**
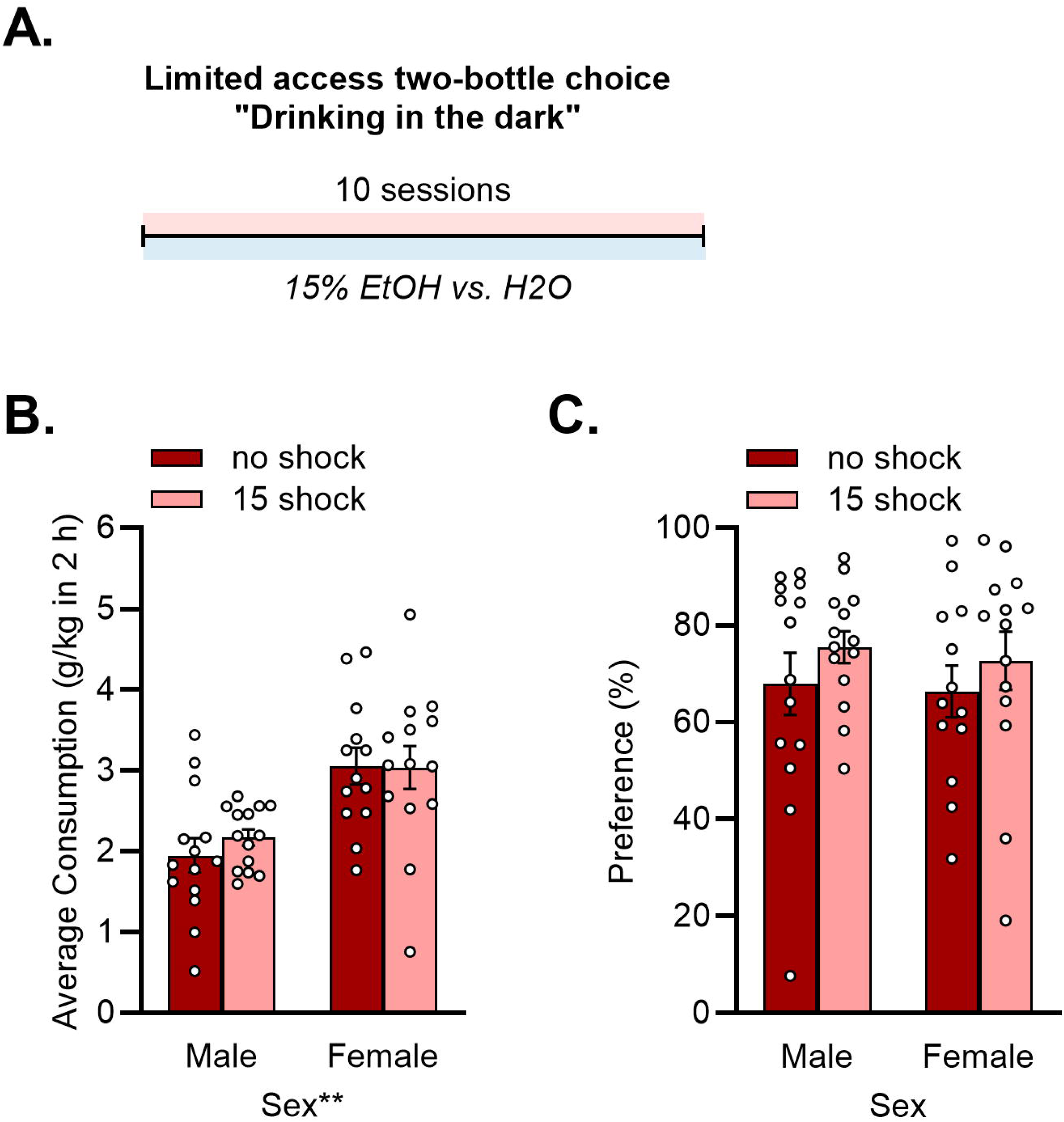
Sex but not infant footshock influences ethanol consumption in a limited access drinking task. (**A**) Mice (n = 13-14/group) drank 15% ethanol for ten 2-h sessions. (**B**) There was a significant influence of sex on ethanol consumption (g/kg), *p < 0.001 main effect of sex. (**C**) Neither sex nor stress altered preference for ethanol vs. water.

#### Experiment 3: Intermittent access ethanol drinking in adulthood

Sixty-six mice (37 male and 24 female) were presented with 20% ethanol following a two-bottle choice, intermittent access protocol for 5 weeks (**Figure 3A**). Ethanol-drinking sessions took place on Monday, Wednesday, and Friday. On intervening sessions (Tuesday, Thursday, Saturday, and Sunday), mice had access to two bottles of drinking water only. Bottle weights were recorded at the beginning of each ethanol-drinking session, after 30 min, and after 24 h. After 4 weeks (12 sessions) of drinking, quinine hemisulfate was added to the ethanol bottle in increasing concentrations of 10 mg/L, 100 mg/L, and 200 mg/L (equivalent to 25, 250, and 500 µM).

**Figure 3.**
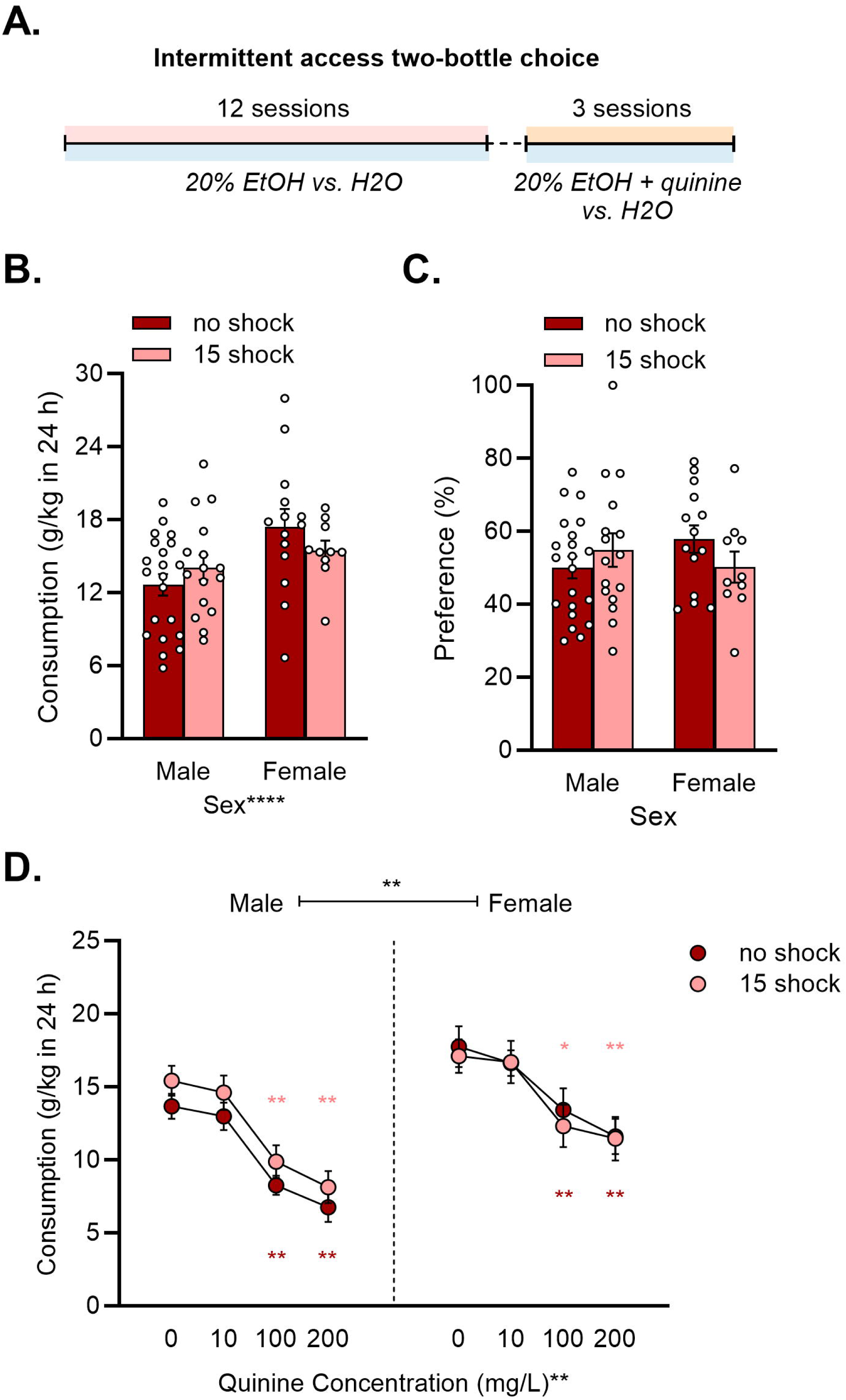
Sex but not infant footshock influences ethanol consumption in an intermittent access drinking task. (**A**) Mice n = 10-21/group) drank 20% ethanol for twelve 24-h sessions. Quinine was added to the ethanol solution on sessions 13-15 in increasing concentrations (10, 100, and 200 mg/L). (**B**) Females consumed more ethanol than males, **p < 0.001 main effect of sex. Infant footshock did not impact ethanol consumption. (**C**) Neither sex nor stress altered preference for ethanol vs. water. (**D**) Females consumed more ethanol mixed with quinine than males, **p < 0.001 main effect of sex. Quinine decreased ethanol consumption regardless of sex or footshock exposure, **p < 0.001 main effect of quinine concentration. Consumption was reduced at the 100 and 200 mg/L concentrations, but not 10 mg/L, in all groups, * p < 0.05, **P < 0.01 Dunnett’s test.

#### Experiment 4: Intermittent access ethanol drinking in adolescence and adulthood

Following ELS on PND 17, 40 mice (21 male and 27 female) underwent a 3-week intermittent access procedure where they received 20% ethanol and RO water on Mondays, Wednesdays, and Fridays between PND 35-55 (**Figure 4A**). On intervening days, mice received two bottles of RO water. Mice were fear conditioned on PND 60 after completing adolescent intermittent access (data not shown). After fear conditioning, mice started adult intermittent access for 5 weeks (17 male and 24 female) (**Figure 5A**). Mice received 20% ethanol and RO water every other day. On intervening days, mice received RO water. During week 5, escalating concentrations of quinine hemisulfate (10 mg/L, 100 mg/L, and 200 mg/L) were mixed with 20% ethanol on Monday, Wednesday, and Friday.

**Figure 4.**
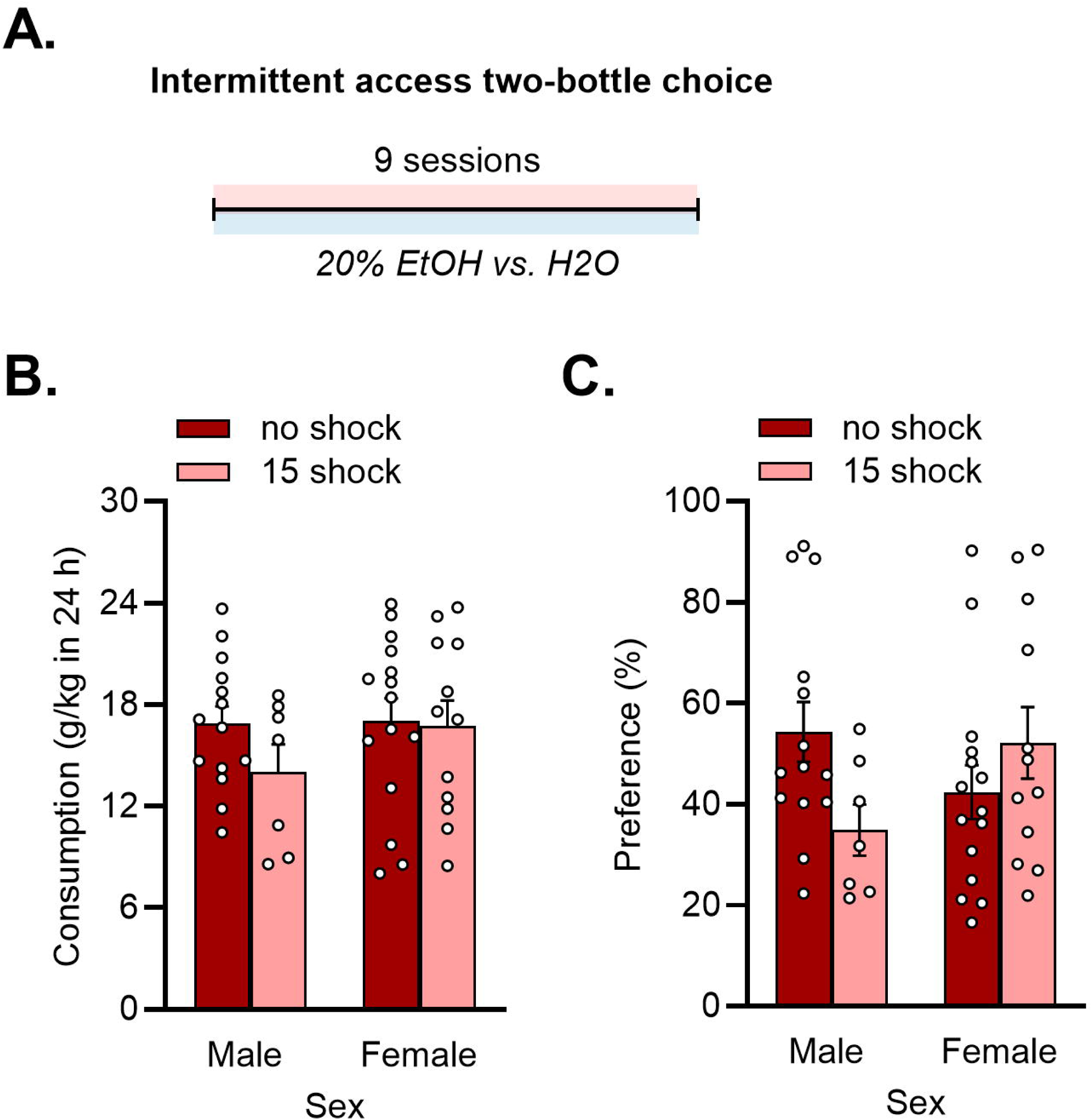
Infant footshock and sex do not influence ethanol consumption in adolescent mice in an intermittent access drinking task. (**A**) Mice (n = 7-14/group) drank 20% ethanol for nine 24-h sessions. (**B**) Infant footshock did not impact ethanol consumption at adolescence. (**C**) Infant footshock did not impact ethanol preference at adolescence.

**Figure 5.**
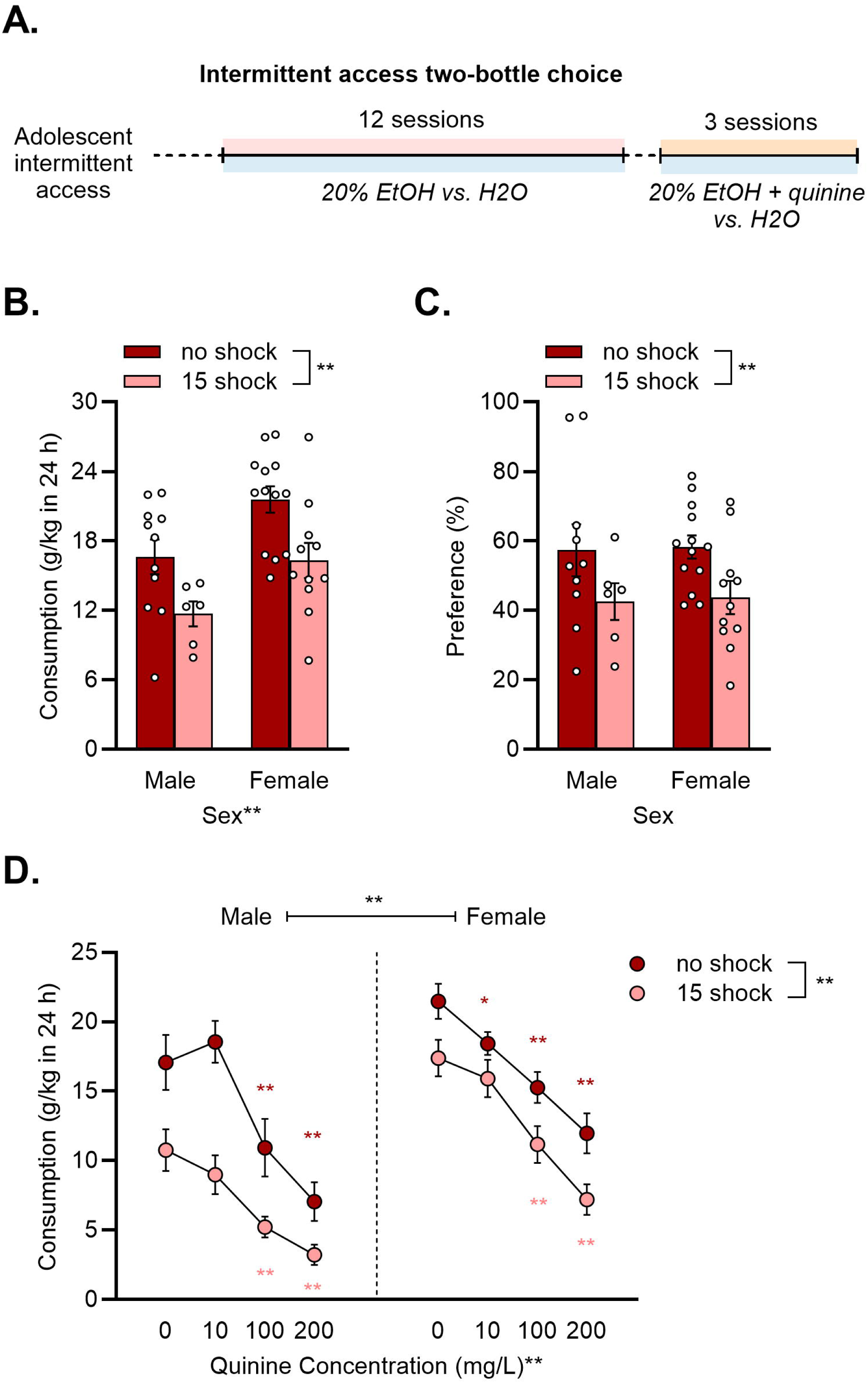
Infant footshock paired with adolescent alcohol exposure influences ethanol consumption and aversion-resistant drinking. (**A**) Mice (n = 6-13/group) drank 20% ethanol for twelve 24-h sessions. Quinine was added to the ethanol solution on sessions 13-15 in increasing concentrations (10, 100, and 200 mg/L). (**B**) 15 shock mice had significantly lower ethanol consumption compared to no shock animals, **p < 0.01 main effect of shock. Female mice drank significantly more ethanol than their male counterparts, **p < 0.01 main effect of sex. (**C**) 15 shock mice had significantly lower ethanol preference compared to no shock animals, *p < 0.05 main effect of shock. (**D**) 15 shock mice had significantly lower ethanol consumption compared to no shock animals, **p < 0.01 main effect of shock. Female mice drank significantly more ethanol than their male counterparts, **p < 0.01 main effect of sex. Quinine decreased ethanol consumption regardless of sex or footshock exposure, **p < 0.01 main effect of quinine concentration. Consumption was reduced at the 100 and 200 mg/L concentrations for all groups, except for no shock females, who also decreased consumption compared to baseline at 10 mg/L, *p < 0.05, **p < 0.01 Dunnett’s test.

Following the fifth week, mice were given only RO water for a week, before undergoing a quinine sensitivity test as a control. For this test, mice were given two bottles of RO water on Monday to establish a baseline. After establishing this baseline, mice were given escalating concentrations of quinine (10 mg/L, 100 mg/L, and 200 mg/L) in one of the RO water bottles on Wednesday, Friday, and the following Monday.

### Data Analysis

Bottle weights were converted to grams (g) of ethanol or milliliters (mL) of water for each mouse. Consumption was calculated as = *(Initial Bottle Weight – Final Bottle Weight) – Average Weight Loss of Dummy Bottles*. Preference was calculated as = *(Volume of Ethanol [Volume of Ethanol + Volume of Water]) x 100*. Consumption and preference were averaged across mice for each group and analyzed using three-way repeated measures (RM) ANOVA with infant footshock, sex, and concentration as factors. All results were first analyzed with session as a factor, but as no significant interaction effects were found, data were averaged across sessions for final analysis and visualization. In the adult intermittent access experiment, social buffering condition (i.e., isolated vs. littermates) was included as a factor, but the data were ultimately collapsed across as there were no significant effects. The Greenhouse-Geisser correction was applied when the assumption of sphericity was violated (ε < 0.75). Follow-up Holm-Sidak’s tests were used as appropriate to assess differences between groups. Aversion-resistant drinking was defined *a priori* as a significant reduction from baseline (= 0 g/L) consumption and was assessed in all groups with a Dunnett’s test to correct for multiple comparisons.

## Results

### Experiment 1: Continuous access ethanol drinking is not altered by infant footshock exposure

Sex, but not stress, impacted ethanol consumption in the continuous access drinking test, with females consuming more than males (**Figure 1B**). A three-way ANOVA revealed no effect of infant footshock on average ethanol consumption across the two concentrations of ethanol presented (F_(1, 36)_ = 0.085, p = 0.773). There was, however, a significant main effect of sex on ethanol consumption (F_(1, 36)_ = 21.300, p < 0.001) and a main effect of ethanol concentration on consumption (F_(1, 36)_ = 10.550, p = 0.003). An interaction between ethanol concentration, sex, and infant footshock neared significance (F_(1, 36)_ = 3.943, p = 0.055). There were no other significant interactions between the factors (all p > 0.409). Holm Sidak’s multiple comparison tests comparing consumption of the 5% and 10% concentrations for each group, demonstrated a greater consumption of 5% vs. 10% ethanol in 15-shock females only (p = 0.008).

Similar to consumption, preference for ethanol vs. water was higher in females than males but was not affected by stress (**Figure 1C**). A three-way ANOVA revealed no significant effect of infant footshock (F_(1, 36)_ = 0.009, p = 0.925) or concentration (F_(1, 36)_ = 3.125, p = 0.086) on ethanol preference. There was, however, a significant main effect of sex on ethanol preference (F_(1, 36)_ = 11.220, p = 0.002). There was also a significant interaction between ethanol concentration, sex, and infant footshock (F_(1, 36)_ = 5.182, p = 0.029). There were no other significant interactions between the factors (all p > 0.487). Holm Sidak’s multiple comparison tests comparing preference for ethanol across concentrations in each group found that 15-shock females had significantly higher preference for 5% vs. 10% ethanol (p = 0.047).

### Experiment 2: Limited access drinking in the dark is not altered by infant footshock exposure

When assessing limited access consumption, there were no effects of stress, but females consumed more than males (**Figure 2B**). A two-way ANOVA revealed no significant effect of infant footshock on ethanol consumption (F_(1, 51)_ = 0.245, p = 0.623). Analysis, however, indicated a significant main effect of sex (F_(1, 51)_ = 22.100, p < 0.001). The interaction between the factors was not significant (F_(1, 51)_ = 0.328, p = 0.569). For ethanol preference, there were no main effects of sex (F_(1, 51)_ = 0.165, p = 0.687) or stress (F_(1, 51)_ = 1.637, p *=* 0.207) and the interaction was not significant (F_(1, 51)_ = 0.012, p = 0.912) (**Figure 2C**).

### Experiment 3: Intermittent access ethanol drinking in adulthood is not altered by infant footshock exposure

We next assessed whether infant footshock influenced adult drinking using an intermittent access drinking task. There were no effects of social buffering observed on any measure, therefore data from mice housed alone vs. with littermates following the ELS session on PND 17 were collapsed for visualization and analysis. Additionally, mice in this experiment received a variable number of footshocks (between 6 and 15) due to an equipment malfunction (average number of shocks males = 11.56 ± 0.72, females = 10.40 ± 0.90) but still displayed significant increases in fear conditioning (data not shown).

Ethanol consumption averaged across the first twelve sessions of intermittent access drinking was higher in females but not affected by stress (**Figure 3B**). A two-way ANOVA revealed no significant effect of infant footshock on average consumption of ethanol (F_(1, 57)_ = 0.057, p = 0.812). There was, however, a significant main effect of sex on average ethanol consumption (F_(1, 57)_ = 7.490, p = 0.008). No significant interaction effect was indicated (F_(1, 57)_ = 2.289, p *=* 0.136). Ethanol preference was also similar across groups, regardless of sex or stress exposure (**Figure 3C**). A two-way ANOVA revealed no significant effect of infant footshock (F_(1, 57)_ = 0.133, p = 0.717), or sex (F_(1, 57)_ = 0.148, p = 0.702) on average preference for ethanol vs. water. No significant interaction effect was indicated, (F_(1, 57)_ = 2.365, p *=* 0.130). To determine whether the absence of a stress effect was due to an insufficient number of shocks in some animals, we correlated the number of shocks received on PND 17 with average consumption (males: r = 0.070, p = 0.796; females: r = -0.227, p = 0.528) and preference (males: r = -0.385, p = 0.131; females: r = -0.096, p = 0.791). These correlations were not significant, further supporting the idea that infant footshock did not influence consumption or preference in this experiment.

For aversion-resistant drinking on sessions 12-15, females continued to consume more than males, and quinine reduced consumption equally in all groups (**Figure 3D**). A three-way ANOVA revealed no significant effect of infant footshock on consumption of quinine adulterated ethanol across sessions 12-15. (F_(1, 57)_ = 0.403, p = 0.528). There were, however, significant main effects of sex (F_(1, 57)_ = 14.590, p < 0.001) and quinine concentration (F_(3, 171)_ = 56.800, p < 0.001) on consumption. No significant interaction effects were indicated (all p > 0.257). To assess aversion resistant consumption, two-way ANOVAs followed by Dunnett’s multiple comparisons tests were performed for each sex. In males, there was a significant main effect of quinine concentration (F_(3, 105)_ = 54.770, p < 0.0001) but no main effect of stress (F_(1, 35)_ = 2.006, p = 0.166) or interaction between the two factors (F_(3, 105)_ = 0.026, p = 0.994).

Compared to baseline, consumption was decreased in no-shock and 15-shock males at the 100 and 200 g/L concentrations (all p < 0.0001), but not at a quinine concentration of 10 g/L (p = 0.770 and 0.754). In females, there was a significant main effect of quinine concentration (F_(3, 66)_ = 14.390, p < 0.0001). Neither the main effect of stress nor interaction reached the threshold for significance. Aversion resistance was observed at the 10 g/L concentration in no-shock (p = 0.802) and 15-shock (p = 0.992) females while consumption was reduced at the 100 and 200 g/L concentrations in both groups (all p < 0.05).

### Experiment 4: Intermittent access ethanol drinking is reduced following exposure to infant footshock and adolescent drinking

Because our prior results demonstrating an effect of infant footshock on drinking in rats included an adolescent drinking period (Radke et al., 2020), we tested the necessity of this exposure in a second cohort of mice that consumed ethanol from PND 35-55. This experiment revealed that infant footshock influences adult consumption when preceded by an adolescent drinking period.

Infant footshock did not affect average ethanol consumption across the nine adolescent drinking sessions (**Figure 4B**). A two-way ANOVA revealed no significant main effects of sex (F_(1, 44)_ = 1.040, p = 0.313), shock (F_(1, 44)_ = 1.223, p = 0.275), or an interaction between sex and shock (F_(1, 44)_ = 0.816, p = 0.371). For average ethanol preference across these sessions, a two-way ANOVA revealed there was a significant interaction between sex and shock (F_(1, 44)_ = 5.126, p < 0.05) (**Figure 4C**). However, there were no significant main effects of sex (F_1, 44)_ = 0.170, p = 0.682) or shock (F_(1, 44)_ = 0.571, p = 0.454) and follow-up Holm-Sidak’s tests comparing preference between shocked groups in males and females were not significant (p > 0.108).

In mice that consumed ethanol during adolescence, infant footshock affected average adult ethanol consumption across twelve sessions (**Figure 5B**). A two-way ANOVA revealed significant main effects of sex (F_(1, 37)_ = 10.870, p < 0.01) and shock (F_(1, 37)_ = 12.250, p < 0.01). However, there was no significant interaction between sex and shock (F_(1, 37)_ = 0.0143, p = 0.906), demonstrating that infant footshock exposure produced lower levels of adult alcohol consumption in mice that drank alcohol as adolescents. For average ethanol preference across twelve adult drinking sessions, a two-way ANOVA revealed a significant main effect of shock (F_(1, 36)_ = 7.006, p < 0.05), but not sex (F_(1, 36)_ = 0.037, p = 0.8498) (**Figure 5C**). Additionally, there was no significant interaction between sex and shock (F_(1, 36)_ = 0.0006, p = 0.981).

Consumption of quinine-adulterated ethanol was affected by infant footshock in mice that consumed ethanol during adolescence (**Figure 5D**). A three-way ANOVA revealed significant main effects of quinine concentration (F_(2.724, 98.05)_ = 81.760, p < 0.0001), sex (F_(1, 36)_ = 14.530, p < 0.001), and shock (F_(1, 36)_ = 17.810, p < 0.001). There was also a significant interaction between quinine concentration, sex, and shock (F_(3, 108)_ = 3.045, p < 0.05). However, there were no other significant interaction effects (all p > 0.3111). To assess aversion-resistant ethanol consumption, two-way ANOVAs followed by Dunnett’s multiple comparisons tests were performed for each sex. In males, there was a significant main effect of quinine concentration (F_(2.096, 29.34)_ = 25.330, p < 0.0001) and shock (F_(1, 14)_ = 10.550, p < 0.01), but no interaction between quinine concentration and shock (F_(3, 42)_ = 1.908, p = 0.143). Compared to baseline, consumption was significantly decreased in both no shock and 15 shock males at the 100 and 200 mg/L concentrations (all < 0.01), but not at 10 mg/L (p = 0.807 and 0.211). In females, there was a significant main effect of quinine concentration (F_(3, 66)_ = 67.770, p < 0.0001) and shock (F_(1, 22)_ = 6.873, p < 0.05), but no interaction between quinine concentration and shock (F_(3, 66)_ = 0.822, p = 0.486). Compared to baseline, consumption was significantly decreased in both no-shock and 15-shock females at the 100 and 200 mg/L concentrations (all p < 0.0001). However, the 15-shock females demonstrated aversion resistance, as they did not significantly decrease consumption compared to baseline at 10 mg/L (p = 0.405), while the no-shock females did decrease (p < 0.05).

As a control, we examined sensitivity to quinine by administering quinine in RO drinking water one week after the cessation of ethanol drinking. This experiment revealed no effects of stress or sex on quinine sensitivity in mice exposed to infant footshock and adolescent ethanol (**Table 1**). A three-way RM ANOVA uncovered a main effect of concentration (F_(3, 108)_ = 21.710, p < 0.0001) and a main effect of sex (F_(1, 36)_ = 6.899, p < 0.05), the latter of which agrees with our prior demonstrations of greater water consumption in female vs. male mice (Sneddon et al., 2023, 2022). There were no other significant main effects of interactions (all p > 0.320). To determine which concentrations of quinine produced aversion resistance, follow up two-way ANOVAs and Dunnett’s tests were performed. In males, there was a significant main effect of concentration (F_(3, 42)_ = 14.850, p < 0.0001) but not stress. Consumption of water adulterated with 10 mg/L quinine was not reduced in either group (no shock: p = 0.585, 15 shock: p = 0.975) while consumption at the 100 mg/L (no shock: p = 0.007, 15 shock: p = 0.047) and 200 mg/L (no shock: p < 0.0001, 15 shock: p = 0.010) concentrations was reduced in all animals. Results in females were similar, with a main effect of concentration (F_(3, 66)_ = 13.980, p < 0.0001) but not stress. Consumption was significantly reduced from baseline at the 100 mg/L (no shock: p = 0.0003, 15 shock: p = 0.003) and 200 mg/L (no shock: p = 0.002, 15 shock: p = 0.003) concentrations in both groups, but not the 10 mg/L concentration.

**Table 1.** Consumption of quinine in water over 24 h. Sensitivity to quinine was not altered by sex or ELS exposure. * p < 0.05, ** p < 0.01 significant main effect. Values in bold typeface are significantly different from baseline (p < 0.05, Dunnett’s test).

| Sex* | ELS | Concentration (mg/L)** | Consumption (mL/kg) |
| --- | --- | --- | --- |
| Male | no shock | 0 | 88.56 ± 10.81 |
|  |  | 10 | 72.71 ± 13.79 |
|  |  | 100 | **40.88 ± 20.20** |
|  |  | 200 | **3.95 ± 0.81** |
|  | 15 shock | 0 | 68.10 ± 12.89 |
|  |  | 10 | 61.63 ± 11.77 |
|  |  | 100 | **20.67 ± 15.26** |
|  |  | 200 | **8.69 ± 4.50** |
| Female | no shock | 0 | 127.22 ± 6.80 |
|  |  | 10 | 97.33 ± 19.72 |
|  |  | 100 | **34.77 ± 19.68** |
|  |  | 200 | **47.45 ± 19.33** |
|  | 15 shock | 0 | 128.00 ± 9.59 |
|  |  | 10 | 112.86 ± 28.20 |
|  |  | 100 | **43.80 ± 25.39** |
|  |  | 200 | **44.55 ± 20.18** |

## Discussion

The central finding of this study is that acute exposure to footshock on PND 17, using a protocol that enhances adult fear learning (Sneddon et al., 2021), affects alcohol drinking behavior in C57BL/6J mice when followed by a period of adolescent intake. Mice exposed to ELS followed by adolescent alcohol had lower alcohol preference and consumption as adults than no-shock mice. In addition, infant stress affected consumption of quinine-adulterated alcohol only in mice that underwent adolescent drinking. In these mice, shocked females did not avoid the lowest quinine concentration while no-shock females did, although we interpret these results with caution as discussed below. There was also a clear effect of sex on drinking behaviors, such that females consumed more alcohol than males when accounting for body weight. However, preference for alcohol vs. water was only higher in females under continuous access conditions.

In a prior study using Long-Evans rats, which investigated alcohol consumption in an intermittent access paradigm following an adolescent drinking period, we demonstrated that PND 17 footshock increased aversion-resistant drinking in male and female subjects but did not alter consumption of unadulterated alcohol (Radke et al., 2020). The current studies were designed to examine whether these effects could be replicated in mice and with other drinking protocols. After failing to find effects of infant footshock on adult consumption using continuous access, limited access, and intermittent access, we tested the necessity of the adolescent drinking period. Our results revealed an effect of infant footshock on alcohol consumption in mice that consumed alcohol in adolescence, which aligns with the original protocol tested in rats (Radke et al., 2020). Thus, we conclude that the infant SEFL protocol can influence adult alcohol drinking behaviors across species and sexes when combined with adolescent alcohol drinking, although the manifestation of the behavioral change differs in rats and mice. For example, infant footshock had no effect on levels of alcohol consumption and preference in rats but decreased both measures of drinking in mice. Further, infant footshock increased the concentration of quinine in alcohol tolerated by rats of both sexes, while the results in mice were less robust. Female mice in the 15-shock group were resistant to 10 mg/L quinine compared to their no-shock counterparts, but this effect should be interpreted with caution considering that mice also did not decrease consumption when this concentration of quinine was added to water. Although we have demonstrated that 10 mg/L quinine in water is avoided by rats and mice previously (Carvour et al., 2024; Radke et al., 2020), since it was not avoided in this experiment we cannot definitively say that 15-shock females were resistant to aversion.

Because we saw increased aversion-resistance when using the same protocol in rats, we expected the infant footshock protocol to similarly enhance vulnerability to alcohol drinking behaviors in mice. Thus, the observation of decreased consumption and preference was surprising and more difficult to reconcile with the hypothesis that ELS enhances vulnerability to alcohol. One possibility is that infant footshock followed by adolescent alcohol consumption produces an anhedonic affective state in adults, leading to reductions in reward-seeking (Hu et al., 2020). This hypothesis could be tested by assessing operant responding for other drugs and natural rewards, intracranial self-stimulation, or social interaction. Another consideration is that adolescent drinking often causes an increase in adult alcohol consumption (Spear, 2016) that may have been prevented in shocked mice. Because Experiments 3 and 4 were conducted separately, we cannot statistically compare the results. However, there is a notable trend toward increased consumption in the no-shock animals between the two experiments (Experiment 3 average consumption male = 12.66 ± 0.89, female = 17.43 ± 1.45; Experiment 4 average consumption male = 16.61 ± 1.50, female = 21.59 ± 1.14) that was not seen in the shocked mice. In either case, our results suggest that ELS and adolescent alcohol consumption engage overlapping mechanisms to produce alterations in adult behavior.

In preclinical studies, stress has been found to have wide-ranging effects on alcohol drinking (e.g., Becker et al., 2011; Radke et al., 2014). At least one prior review article estimated that acute stress increased drinking in fewer than 50% of studies (Becker et al., 2011). The limited and/or variable effects of acute stress exposure on drinking are in contrast with chronic stress exposure, which increase drinking behaviors much more consistently, especially when administered in the early stages of development (Becker et al., 2011; Radke et al., 2014). Further, adolescent alcohol exposure has previously been shown to render adult rats more susceptible to the effects of stressors, including footshock (Füllgrabe et al., 2007; Siegmund et al., 2005). Thus, when considered alongside the literature, the current studies highlight the variability inherent to acute stress effects on alcohol drinking behaviors and the importance of other developmental experiences, be it stress or drug exposure, to adult outcomes. It is important to consider that consumption of alcohol could be described as a stressor in and of itself. Alcohol exposure increased activity in the hypothalamic-pituitary-adrenal (HPA) axis, and withdrawal from alcohol increased levels of corticotropin-releasing factor (Olive et al., 2002; Zorrilla et al., 2001). The intermittent access drinking procedure also requires animals to be individually housed, which is another significant source of stress (Butler et al., 2016; Walker et al., 2019). Alcohol drinking in adolescence may consequently reveal the effects of infant footshock on adult drinking by acting as a secondary stressor that potentiates the effects of ELS. Further studies will be necessary to determine whether alcohol exposure alone, without the stress of individual housing, can replicate the effects seen here and in our rat studies.

Our final experiment incorporated additional stressors beyond adolescent alcohol exposure. Mice in this study were isolated from their dam and littermates following footshock and were fear-conditioned during adulthood. Based on the results of Experiment 3, and our studies in rats, isolation from littermates does not appear to be a necessary condition to produce effects on drinking, although it is unclear at present whether isolation from the dam contributes to the ELS effects. Adult fear conditioning was not incorporated in the drinking in the dark or continuous access experiments, which could possibly drive the absence of a drinking effect in those studies. However, the lack of an effect in Experiment 3, which did include adult fear conditioning, makes it more likely that adolescent drinking is the key factor. These factors could be tested more systematically in the future to determine the number of stressful experiences necessary to produce an effect on alcohol drinking behaviors.

It is important to consider why the effects of infant stress followed by adolescent alcohol drinking were model-species dependent. There are several notable differences between Long-Evans rats and C57BL/6J mice. Long-Evans rats are an outbred line, and thus in a given set of subjects there is increased genetic diversity compared to the inbred C57BL/6J line. The effect of stress on alcohol consumption has been found to vary between mouse strains (Radke et al., 2014). Furthermore, PTSD-AUD comorbidity is not ubiquitous in individuals who have experienced trauma (Gilpin and Weiner, 2017; Young-Wolff et al., 2011). Thus, the effect of early life trauma is likely dependent on genetic predispositions (Koenen et al., 2009; Young-Wolff et al., 2011). Another notable difference between C57BL/6J mice and Long-Evans rats is that C57BL/6J mice are bred to be high sucrose-preferring mice and are known to consume more alcohol than other strains of mice. In contrast, Long-Evans rats have no predisposition to maladaptive alcohol drinking. Thus, consumption may not be high enough in Long-Evans rats to observe reductions in drinking like those seen in mice.

Our results show a clear effect of sex on alcohol consumption, with females consuming more alcohol than males, which is consistent with a large body of previous literature (Finn, 2020; Radke et al., 2021a, 2021b). Further, males and females displayed similar preference for alcohol vs. water in the limited access and intermittent access experiments. This finding is consistent with previous studies of sex differences in alcohol drinking behaviors in our lab that have found divergences in the effects of biological sex on alcohol consumption vs. preference (Sneddon et al., 2022, 2019), and may give insight into the origin of sex differences in alcohol drinking. Sex differences in alcohol preference have been found to differ based on alcohol concentration (Radke et al., 2021b; Sneddon et al., 2022), but it is unclear if the lack of an effect on preference seen here resulted from the different concentrations of alcohol used in the three experiments or from the intermittent access vs. continuous access conditions. It is also interesting that female rats are more sensitive to the effects of infant footshock on adult fear behaviors (Minshall et al., 2024) than males and that shocked females demonstrated greater aversion-resistance in the studies reported here. Thus, the model developed here may capture the increased vulnerability for comorbid PTSD-AUD observed in female patients (Breslau et al., 1997; Gavranidou and Rosner, 2003).

In summary, our study found no effect of infant footshock on alcohol drinking behaviors in mice unless a period of adolescent alcohol drinking was included. A major goal of this line of work is to establish a viable model of comorbid PTSD and AUD in male and female rodents. Infant footshock is appealing for its relatively selective effects on PTSD-relevant behaviors, including fear learning and anxiety behaviors (Poulos et al., 2014; Quinn et al., 2014) as well as resistance to fear extinction (Sneddon et al., 2021). This model is also unique in that stress is administered prior to weaning and before animals develop the ability to form associative memories, unlike other models in which stress exposure occurs during adolescence (Shaw et al., 2020; Skelly et al., 2015). Importantly, the current model has also been validated in male and female animals. It should be noted that contradictory results and poor replicability are common in studies investigating the relationship between stress and alcohol exposure. Thus, in order to understand the intricate differences in the relationship between stressors and alcohol consumption, future studies should continue to investigate the specific conditions necessary to increase alcohol drinking following ELS and include measurements of physical indicators of the stress response such as corticosterone levels. Such continued development and refinement of mouse models of comorbid PTSD and AUD will be beneficial to uncovering the shared neural mechanisms of these two debilitating disorders.

## Acknowledgements

The authors are grateful to Caroline Scribner, Brandon Arnold, and Annemarie Thomas for assistance with behavioral experiments.

## Author contributions

Thomas Perry: Formal Analysis, Investigation, Visualization, Writing – Original Draft. Harrison Carvour: Formal Analysis, Investigation, Visualization, Writing - Original Draft. Amanda Reichert: Formal Analysis, Investigation, Writing – Original Draft. Elizabeth Sneddon: Formal Analysis, Investigation, Writing – Original Draft. Charlotte Roemer: Formal Analysis, Investigation. Ying Ying Gao: Formal Analysis, Investigation. Kristen Schuh: Formal Analysis, Investigation. Natalie Shand: Investigation, Writing – Original Draft. Jennifer Quinn: Conceptualization, Funding Acquisition, Supervision, Writing - Review and Editing. Anna Radke: Conceptualization, Funding Acquisition, Supervision, Visualization, Writing - Review and Editing.

